# Evidence for non-methanogenic metabolisms in globally distributed archaeal clades basal to the *Methanomassiliicoccales*

**DOI:** 10.1101/2020.03.09.984617

**Authors:** Laura A. Zinke, Paul N. Evans, Alena L. Schroeder, Donovan H. Parks, Ruth K. Varner, Virginia I. Rich, Gene W. Tyson, Joanne B. Emerson

## Abstract

Recent discoveries of *mcr* and *mcr*-like complexes in genomes from diverse archaeal lineages suggest that methane (and more broadly alkane) metabolism is an ancient pathway with complicated evolutionary histories. The conventional view is that methanogenesis is an ancestral metabolism of the archaeal class *Thermoplasmata*. Through comparative genomic analysis of 12 *Thermoplasmata* metagenome-assembled genomes (MAGs), we show that these microorganisms do not encode the genes required for methanogenesis, which suggests that this metabolism may have been laterally acquired by an ancestor of the order *Methanomassiliicoccales*. These MAGs include representatives from four orders basal to the *Methanomassiliicoccales*, including a high-quality MAG (95% complete) that likely represents a new order, *Ca.* Lunaplasma lacustris ord. nov. sp. nov. These MAGs are predicted to use diverse energy conservation pathways, such as heterotrophy, sulfur and hydrogen metabolism, denitrification, and fermentation. Two of these lineages are globally widespread among anoxic, sedimentary environments, with the exception of *Ca.* Lunaplasma lacustris, which has thus far only been detected in alpine caves and subarctic lake sediments. These findings advance our understanding of the metabolic potential, ecology, and global distribution of the *Thermoplasmata* and provide new insights into the evolutionary history of methanogenesis within the *Thermoplasmata*.

## Introduction

High-throughput sequencing of environmental DNA, metagenomic assembly of DNA sequence reads into contigs, and binning of contigs into metagenome-assembled genomes (MAGs) has provided unprecedented insights into the metabolic potential and evolutionary history of many uncultivated lineages (Hug *et al.*, 2016; Brown *et al.*, 2015; Woodcroft *et al.*, 2018; Castelle *et al.*, 2013; Crits-Christoph *et al.*, 2018; Castelle *et al.*, 2015; Anantharaman *et al.*, 2016). In addition to revealing numerous new phyla, MAG analyses have identified new clades of microorganisms within longstanding phylogenetic groups, assigned biogeochemical roles to known but uncultivated lineages, and attributed new functions to diverse relatives of model prokaryotes (Graham *et al.*, 2018; Mondav *et al.*, 2014; Tully, 2019; Boyd *et al.*, 2019; Solden *et al.*, 2016; Singleton *et al.*, 2018; Martinez *et al.*, 2019). With these advances in sequencing technology, a more complex view of microbial evolution and metabolism has emerged. Increasingly, metabolic divergence between even closely related organisms is now recognized, with suggestions that some metabolism types, such as denitrification, are readily and repeatedly transferred to other microorganisms (Meyer and Kuever, 2007).

Within the last decade, our view of alkane/methane metabolism has evolved from being ascribed to specific *Euryarchaeota* orders to being found in multiple archaeal phyla (Martiny *et al.*, 2015; Evans *et al.*, 2019; Vanwonterghem *et al.*, 2016; Borrel *et al.*, 2019). In addition to the phylogenetic diversity of methanogens, more variations on the methane-cycling biochemical pathways have been discovered (as in Evans *et al.*, 2019; Hua *et al.*, 2019). Genes typically associated with methane metabolism have also been linked to alkane oxidation in multiple Archaea (Laso-Pérez *et al.*). These findings have led to vigorous investigation into the evolutionary history of methanogens and the Archaea (e.g. Wolfe and Fournier, 2018a; Roger and Susko, 2018; Wolfe and Fournier, 2018b; Berghuis *et al.*, 2019; Spang *et al.*, 2017; Borrel *et al.*, 2019). For example, it is hypothesized that the short H_2_-dependent methylotrophic methanogenesis pathway might be transferred through horizontal gene transfer more easily than the longer H_2_/CO_2_ dependent pathway (Borrel *et al.*, 2016; Evans *et al.*, 2019), though a consensus has not been reached.

The current understanding of methanogenesis evolution is that ancestors of the *Euryarchaeota* phylum were methanogens (Evans *et al.*, 2019), or potentially methanotrophs or alkanotrophs (Spang *et al.*, 2017), with methanotrophy-driven acetogenesis also being proposed as an early metabolism type (Russell and Nitschke, 2017). It has been suggested that some euryarchaeal lineages, such as the *Thermoplasmatales*, lost the methanogenesis pathway, while others, such as the *Methanomassiliicoccales*, retained at least part of the pathway (Evans *et al.*, 2019). However, a recent taxonomic reclassification based on relative evolutionary distance (RED) has proposed a new phylum, *Ca.* Thermoplasmatota, which includes the *Thermoplasmatales*, the *Methanomassiliicoccales*, the *Aciduliprofundales*, the MGII/MGIII archaea, and other uncharacterized lineages (Rinke *et al.*, 2019). Within *Ca.* Thermoplasmatota, only the *Methanomassiliicoccales* are known methanogens (Borrel *et al.*, 2014; Dridi *et al.*, 2012), using a truncated H_2_-dependent methylotrophic methanogenesis pathway (Lang *et al.*, 2015), while the other *Ca.* Thermoplasmatota lineages utilize a diverse set of metabolic strategies for energy conservation (Rinke *et al.*, 2019; Tully, 2019; Sapra *et al.*, 2003). This suggests that the basal ancestor to the *Ca.* Thermoplasmatota potentially was not a methanogen, which this study further supports.

Here, MAGs from several clades that are phylogenetically close to the methylotrophic methanogenic *Methanomassiliicoccales* are described. These lineages are consistently basal to the *Methanomassiliicoccales* in phylogenetic trees, although in some cases these MAGs were previously classified as *Methanomassiliicoccales* in public databases. Phylogenetic trees, MAG annotation, and functional profiling revealed key differences between these clades and *Methanomassiliicoccales*, most notably a lack of methanogenesis potential in the basal clades. Finally, the global distribution of three of these populations was determined and a new order within the *Thermoplasmata*, sister to the *Methanomassiliicoccales* and the *Thermoplasmatales,* is proposed, named here as *Ca*. Lunaplasmatales ord. nov.

## Materials and Methods

### Data acquisition

Previous work suggested that a MAG, referred to here as subarctic lake (SAL) 16, assembled from methanogenic sediments of a subarctic lake in northern Sweden, lacked methanogenesis genes, despite being assigned as a *Methanomassiliicoccales* based on the genomic information and RDP assignment of the assembled and binned 16S rRNA gene (Seitz *et al.*, 2016). To further investigate whether these missing genes were the result of incomplete genome binning or were likely to be a true lack of methanogenic capabilities, MAGs were identified that were closely related to SAL16 in the Genome Taxonomy Database (GDTB; https://gtdb.ecogenomic.org/tree) release 86 archaeal tree, including those which were also in the *Ca.* Thermoplasmata_A class in GTDB (Parks *et al.*, 2018). Additionally, publicly available MAGs and genomes within the *Thermoplasmata* and *Ca.* Thermoplasmata_A classes, which together encompass the *Methanomassiliicoccales*, *Thermoplasmatales*, and *Aciduliprofundum* (Parks *et al.*, 2018), were included here for comparison. MAGs and genomes were downloaded from the National Center for Biotechnology Information (NCBI) database from the accession numbers, the GTDB, or from the publications listed in Supplemental Table 1. MAGs beginning with the “UBA” prefix have been binned and published previously as described in Parks *et al.*, 2017, while those beginning with “RBG” prefix are from Anantharaman *et al.*, 2016, SAL16 is from Emerson *et al*. 2020, and SG8-5 is from Lazar *et al.*, 2017. Genome statistics for all MAGs/genomes were determined using CheckM (Parks *et al.*, 2015; Supplemental Table 2). The completeness and redundancy of the 12 MAGs of interest were further assessed using the MiGA webserver (http://enve-omics.ce.gatech.edu:3000/, accessed November, 2018; Rodriguez-R *et al.*, 2018; Supplemental Table 3).

### MAG identification

MAGs were screened for 16S rRNA gene sequences using the MiGA webserver (Rodriguez-R *et al.*, 2018). Three MAGs contained 16S rRNA gene sequences, which were classified using the Ribosomal Database Project (RDP) classifier (Wang *et al.*, 2007). The 16S rRNA gene sequences were also uploaded to the SILVA Alignment, Classification and Tree (ACT) service (Quast *et al.*, 2012; Pruesse *et al.*, 2007). The sequences were aligned to the global SILVA SSU alignment and classified with a minimum identity of 0.95 and 10 neighbor sequences (Pruesse *et al.*, 2012). The 16S rRNA gene sequences were also compared to the NCBI nucleotide (nt) database using BLASTn for additional insight into taxonomy.

The 12 MAGs of interest were additionally compared to the NCBI Genome database (prokaryotes) using the MiGA server (Rodriguez-R *et al.*, 2018) (Supplemental Tables 5 and 6). Genome and MAG taxonomy was also compared to GTDB taxonomy, which uses RED values to determine taxonomic groupings (Parks *et al.*, 2018). The GTDB Toolkit (GTDB-Tk) v0.1.3 classification workflow (https://github.com/Ecogenomics/GtdbTk) was used to determine RED-based taxonomic placement of MAGs not already in the GTDB database (http://gtdb.ecogenomic.org/) (Supplemental Table 5). Pairwise average amino acid identities (AAIs) between all genomes and MAGs were computed using the envi-omics AAI calculator (http://enve-omics.ce.gatech.edu/aai/<u>, accessed January 2019</u>; Rodriguez-R and Konstantinidis, 2016) (Supplemental Table 7).

### Phylogenetic trees

The phylogenetic tree based on 16 concatenated ribosomal proteins used in (Hug *et al.*, 2016) (Figure 1) was constructed using the methodology outlined in (Graham *et al.*, 2018). Briefly, open reading frames in MAGs and genomes were determined using Prodigal v2.6.3 (Hyatt *et al.*, 2010), and ribosomal proteins were identified from these ORFs with HMMER v3.1b2 using the command hmmsearch with an e-value cutoff of 1E-5 (Eddy, 2011). Individual proteins were aligned in Muscle v3.8.31 (Edgar, 2004), and alignments were trimmed using TrimAL v.1.2rev59 in automatic1 mode (Capella-Gutierrez *et al.*, 2009). Proteins were concatenated, and a maximum likelihood tree was calculated using FastTree v.2.1.10 with the parameters -lg -gamma and 1000 bootstraps (Price *et al.*, 2010; 2009). The resulting tree was visualized through the interactive Tree of Life (iTOL) webserver (Letunic and Bork, 2016).

**Figure 1.**
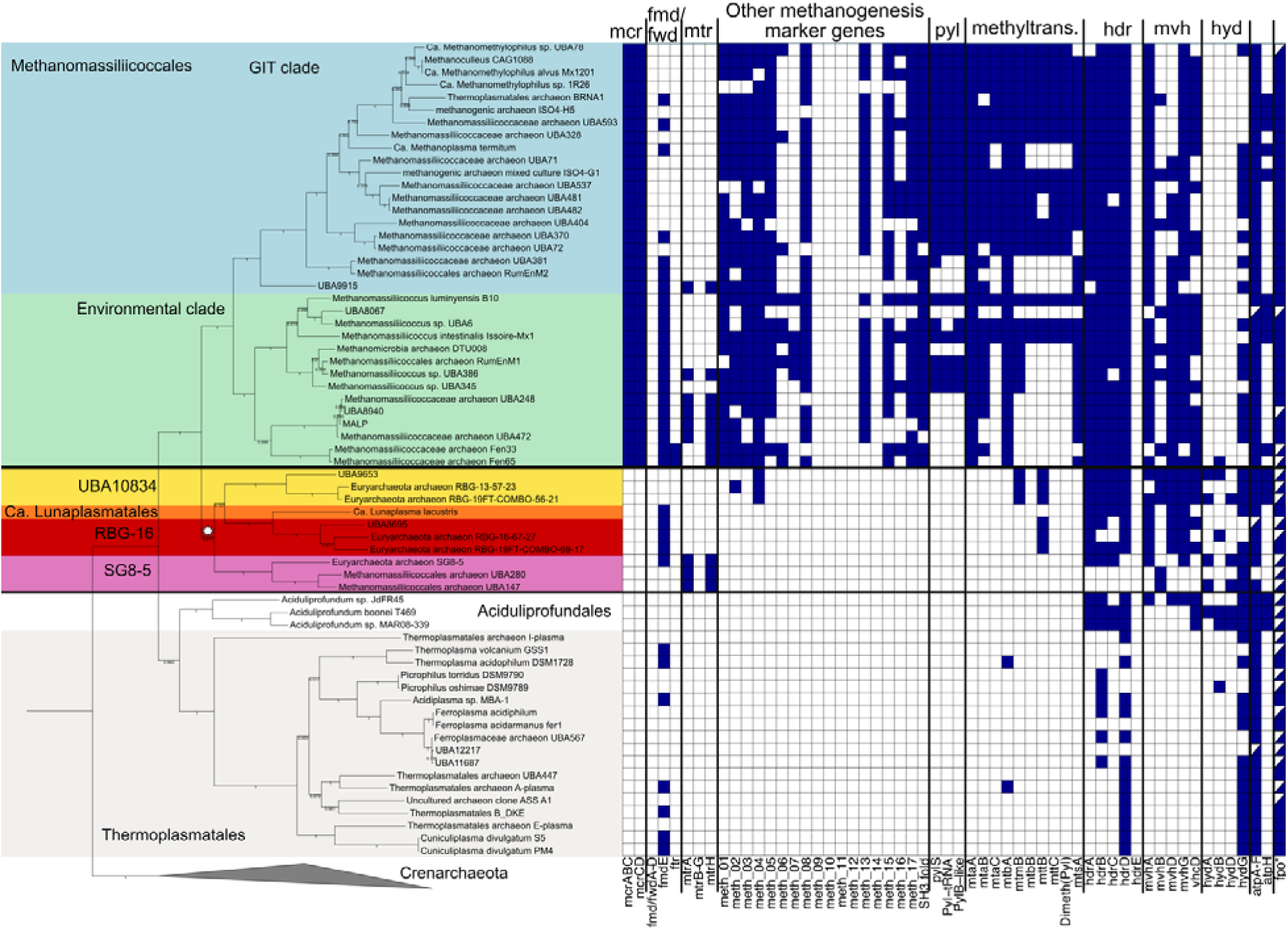
Ribosomal protein (RP) tree. Tree (based on the 16 proteins used in Hug *et al*., 2016), showing the phylogenetic relationships among 10 of the non-methanogenic MAGs basal to the *Methanomassiliicoccales* (in gold, orange, red, and pink backgrounds) and closely related Archaea, including the *Methanomassiliicoccales* (gastrointestinal tract (GIT) and environmental clades in blue and green backgrounds, respectively, following clade designations from Söllinger *et al.* (2016)), *Aciduliprofundales* (white background), and *Thermoplasmatales* (grey background). The white star represents the proposed gain of methanogenesis in the *Methanomassiliicoccales*. Bootstrap values (of 1000 bootstraps) are shown as a filled black dot where support was higher than 80%, as an empty dot for more than 50% support, and as values for branches with less than 50% support. The heatmap to the right demonstrates the genomic potential for methanogenesis in each of the genomes and MAGs (genes colored blue were detected, white were not, half-filled boxes indicate where some subunits were missing; see Supplemental Table 8 for further gene information). Methanogenesis marker genes are those listed in Borrel *et al*. 2014. Two MAGs in the UBA10834 group are not included in the tree because they lack at least 8 of the 16 RP required to be included in the tree. The methanogenesis complement for these MAGs is in Supplemental Figure 5. *Fpo includes subunits ABCDHIJKLMN.

The 16S rRNA gene tree (Supplemental Figure 1) was made using full-length or near full-length sequences from the 14 MAGs and genomes that had 16S rRNA sequences (*Thermoplasmatales* archaeon BRNA1, *Methanomassiliicoccus* sp. UBA386, *Methanomassiliicoccus* sp. UBA345, *Methanomassiliicoccus* sp. UBA6, *Methanomassiliicoccaceae* archaeon UBA593, *Ca.* Methanomethylophilus alvus Mx1201, methanogenic archaeon ISO4-H5, *Aciduliprofundum boonei* T469, *Aciduliprofundum* sp. MAR08-339, *Aciduliprofundum* sp. EPR07-39, *Thermoplasma acidophilum*, *Cuniculiplasma divulgatum* S5, *Cuniculiplamsa divulgatum* PM4, and *Acidiplasma* sp. MBA-1), as well as sequences from the NCBI database (accessed December, 2018). Sequences were aligned and trimmed using Muscle v3.8.45 implemented in Geneious v11.0.5 (100 maximum iterations) (Edgar, 2004), and a tree was built with RAxML 8.2.12 with the parameters -m GTRCAT -f a -x 123 -p 456 and 1000 bootstraps (Stamatakis, 2014).

The RED-based phylogenetic tree (Supplemental Figure 2) was constructed using the GTDB Toolkit (GTDB-Tk) de_novo workflow (https://github.com/Ecogenomics/GtdbTk), including the GTDB classes Thermoplasmata and Thermoplasmata_A and using Crenarchaeota as the outgroup.

MtrH and MttB trees (Supplemental Figures 3 and 4 and) were made using sequences identified by Prokka v1.13.3, BlastKoala v2.1, and InterProScan v5.30-69.0 in the MAGs and genomes, and select sequences from the NCBI GenBank and UniProt databases (accessed January, 2019). MtrH was selected instead of MtrA for phylogenetic tree reconstruction since more of the *Methanomassiliicoccales* MAGs contained MtrH than MtrA. Sequences were aligned and trimmed using Muscle (Edgar, 2004) implemented in Geneious (100 maximum iterations), and a tree was built with RAxML with the parameters -f a -m PROTGAMMAAUTO -p 12345 -x 12345 and 100 bootstraps (Stamatakis, 2014).

### Functional annotation

For all genomes, putative open reading frames (ORFs) were called, translated to amino acid sequences, and functionally annotated in Prokka with the kingdom set to Archaea (Seemann, 2014). InterProScan was used with default settings to compare the ORF amino acid sequence outputs from Prokka to the InterPro, TIGRFAM, and PFAM databases (Jones *et al.*, 2014). ORFs from the 12 MAGs of interest were also uploaded to the blastKOALA server and annotated with KEGG Orthologies using the Archaea taxonomy setting and the “family_eukaryotes + genus_prokaryotes” KEGG GENES database (Kanehisa *et al.*, 2016) (Supplemental Tables 9-20). These ORFs were also screened to determine which contained export signals using psortb in Archaea mode (Yu *et al.*, 2010). Additional verification through BLASTp against the NCBI nr database was performed for manual curation of MAGs.

### Recovering global 16S rRNA gene sequence distributions

The 16S rRNA gene sequences recovered from SG8-5, UBA147, and SAL16 (*Ca*. Lunaplasma lacustris) (16S rRNA gene sequences were recovered from three of the 12 MAGs) were compared to nucleotide sequences in the NCBI-nt database using methods similar to (Mondav *et al.*, 2014). Briefly, for each 16S rRNA gene sequence, standalone blast v2.2.31 command blastall was used with the settings -v 200000 -b 200000 -p blastn -m 8 against the nt database (downloaded in November, 2018) (Lipman *et al.*, 1997). The results were parsed using the bio-table command (https://github.com/pjotrp/bioruby-table) at 97% similarity and requiring the matched sequence to be at least 200 bp long, which removed spurious hits to conserved regions. Hits were manually searched by their GenBank identifiers. Those that could be associated with peer-reviewed publications and included latitude and longitude of the sample origin were used for mapping the distribution of these organisms (Supplemental Table 21). Additionally, the map included locations of the metagenomes from which the MAGs of interest were recovered (Table 1). The map was created in R using the ‘maps’ and ‘ggplot2’ packages (Brownrigg *et al.*; Wickham, 2011).

**Table 1.** Genome statistics of MAGs from *Ca*. Lunaplasma lacustris and the orders RBG-16, UBA10834, and SG8-5. Completeness, contamination, contigs, and genome size (bp) were produced using CheckM. Sample type, metagenome origin, and reference were retrieved from NCBI or GTDB.

| MAG name | Completeness | Contamination | Contigs | Genome Size | Sample type | Metagenome origin | Reference |
| --- | --- | --- | --- | --- | --- | --- | --- |
| *Ca. Lunaplasma lacustris* | 95.2 | 4.8 | 91 | 2535897 | sediment | Inre and Mellestra Harsjon, Sweden | Emerson et al. *submitted* (on bioRxiv) |
| Euryarchaeota archaeon RBG_19FT_COMBO_69_17 | 96.67 | 6.34 | 449 | 2210690 | sediment | Rifle, Colorado, USA | Anantharaman et al. (2016) Nat. Comms. |
| Euryarchaeota archaeon RBG_19FT_COMBO_56_21 | 92.8 | 0.8 | 110 | 1868766 | sediment | Rifle, Colorado, USA | Anantharaman et al. (2016) Nat. Comms. |
| UBA10834 | 89.79 | 2.4 | 319 | 1705491 | sediment | Noosa River estuary, SE Queensland, Australia | Parks et al. (2017) Nat. Micro. |
| Methanomassiliococcales archaeon UBA147 | 89.6 | 3.2 | 80 | 1995084 | waste water | Suncor tailings pond 6, Alberta, Canada | Parks et al. (2017) Nat. Micro. |
| UBA9653 | 88.67 | 0.8 | 500 | 1711095 | ground water | Rifle, Colorado, USA | Anantharaman et al. (2016) Nat. Comms. |
| Euryarchaeota archaeon RBG_13_57_23 | 83.8 | 0.8 | 108 | 1520809 | sediment | Rifle, Colorado, USA | Anantharaman et al. (2016) Nat. Comms. |
| Euryarchaeota archaeon RBG_16_67_27 | 77.47 | 0.4 | 144 | 1546722 | sediment | Rifle, Colorado, USA | Anantharaman et al. (2016) Nat. Comms. |
| Methanomassiliococcales archaeon UBA280 | 76.8 | 2.4 | 46 | 1749312 | waste water | Suncor tailing pond 5, Alberta, Canada | Parks et al. (2017) Nat. Micro. |
| Euryarchaeota archaeon RBG_16_62_10 | 74.45 | 1.6 | 971 | 1494907 | sediment | Rifle, Colorado, USA | Anantharaman et al. (2016) Nat. Comms. |
| Euryarchaeota archaeon SG8-5 | 72.24 | 0.8 | 114 | 1247229 | sediment | White Oak Estuary, North Carolina, USA | Lazar et al. (2017) ISME J. |
| UBA8695 | 70.76 | 0 | 317 | 1709476 | soil | Murray Bridge, South Australia, Australia | Parks et al. (2017) Nat. Micro. |

## Results and Discussion

### Phylogeny of MAGs within and basal to Methanomassiliicoccales

In our previous work, a MAG recovered from subarctic lake sediment metagenomes in Northern Sweden, referred to here as SAL16, was characterized taxonomically as a member of the archaeal order *Methanomassiliicoccales* (Seitz *et al.*, 2016). However, despite being 95% complete, the MAG lacked genetic evidence for methanogenesis, which was unusual, given that all previously identified *Methanomassiliicoccales* were thought to be methanogens (Borrel *et al.*, 2014; Söllinger *et al.*, 2015). Similarly, study of archaeal genomes recovered from aquifer water and sediments near the Colorado River in Rifle, Colorado suggested that *mcrA* gene sequences (indicative of methanogenesis) were not co-binned with the *Methanomassiliicoccales* or their close relatives, but these *Methanomassiliicoccales-*related MAGs were less than 50% complete, so the authors could not conclude whether or not these organisms were methanogens (Castelle *et al.*, 2015). In order to more fully assess the phylogeny and metabolic potential of the SAL16 MAG and its close relatives, we downloaded MAGs and genomes at least 70% complete from NCBI and the GTDB, targeting those classified as or grouped phylogenetically closely to the *Methanomassiliicoccales*, as well as sister clades *Thermoplasmatales* and *Aciduliprofundales* (GTDB classes Thermoplasmata and Thermoplasmata_A).

In total, 68 MAGs and genomes from these lineages were examined, with a focus on 12 publicly available MAGs (including SAL16) that we found to be closely related to, but not within, the *Methanomassiliicoccales* (Table 1; for MAG accession numbers, see Supplemental Table 1). It is important to note, however, that some of these MAGs were labeled as “Methanomassiliicoccales archaeon” in the NCBI database and categorized as “Methanomassiliicoccus” by the RDP classification tool, while being classified within a separate order or even class by other pipelines, such as GTDB-Tk (Parks *et al.*, 2018), MiGA (Rodriguez-R *et al.*, 2018), SILVA SINA (Pruesse *et al.*, 2007; Yilmaz *et al.*, 2013; Quast *et al.*, 2012), and amino acid identity (AAI) (Rodriguez-R and Konstantinidis, 2016), as described in further detail below. The MAGs range in completeness from 70.8 to 95.2% with up to 6.3% redundancy (Table 1; Supplemental Table 2), as determined by CheckM (Parks *et al.*, 2015). SAL16 is considered a high-quality draft genome by the MIMAG standards (Bowers *et al.*), except for the lack of a binned 5S SSU rRNA gene. SAL16 was characterized as an ‘excellent’ quality genome through the MiGA webserver (Rodriguez-R *et al.*, 2018), with the other 11 MAGs generally categorized as ‘high’ (8 MAGs) or ‘intermediate’ (2 MAGs) quality, apart from one ‘low’ quality MAG (UBA10834) (Supplemental Table 3).

An assembled 16S rRNA gene sequence was recovered from each of three MAGs: SAL16 (1,465 bp), UBA147 (1,174 bp), and SG8-5 (1,459 bp), the last of which was previously determined to group phylogenetically within the Rice Cluster III lineage (Lazar *et al.*, 2017) (Supplemental Figure 1). The SAL16 16S rRNA gene sequence shares 87% nucleotide identity with both the UBA147 and SG8-5 sequences, indicating that these MAGs belong to different orders within the same class. SG8-5 and UBA147 share 95% similarity between their 16S rRNA gene sequences, revealing that they could represent the same genus, though the 59.7% AAI between them only supports that they represent the same family (Konstantinidis *et al.*, 2017). In all cases, the closest cultured relative was *Methanomassiliicoccales luminyensis* B10, at 86% rRNA gene identity with SAL16 and 88-89% with UBA147 and SG8-5, placing these MAGs in at least novel orders within the *Thermoplasmata* class (Supplemental Table 4) (Konstantinidis *et al.*, 2017). In contrast, the Ribosomal Database Project (RDP) classifier (https://rdp.cme.msu.edu/classifier/classifier.jsp) run with a 95% confidence threshold classified all three MAGs within the *Methanomassiliicoccus* genus (Cole *et al.*, 2009) (Supplemental Table 4). If these sequences had been recovered from 16S rRNA-based amplicon studies, they likely would have been assigned as *Methanomassilicoccales*, which could lead to spurious assignments of metabolic capability (*i.e*., an assumption that these sequences represent methanogens).

The RED calculated by the GTDB-Tk workflow (Parks *et al.*, 2018) supported the placement of SAL16 as a novel order within the Thermoplasmata class, which also contains the *Methanomassiliicoccales* (Supplemental Table 5). The 11 other MAGs were previously classified within orders containing no isolated representatives: the orders RBG-16-68-12 (3 MAGs), UBA10834 (5 MAGs), and SG8-5 (3 MAGs) (Supplemental Table 5; Parks *et al.*, 2018). Similarly, taxonomic classification and novelty calculations through the MiGA webserver also indicated that most of these MAGs represent novel orders, and potentially novel classes in some cases (Supplemental Table 5). The five MAGs in the order UBA10834 (RBG_COMBO_56_21; RBG_13_57_23; RBG_16_62_10, UBA10834, and UBA9653 (Figure 1; Supplemental Figure 2; Supplemental Table 5) were estimated to be within the *Thermoplasmata* class, but likely not within the *Methanomassiliicoccales* (Supplemental Table 5). The AAI values between these five MAGs and *Methanomassiliicoccus luminyensis* B10 were 46.9-47.7% AAI (Supplemental Tables 6 and 7), placing them at the lower end of the range suggested for being within the same family (Konstantinidis *et al.*, 2017). These results, taken together with the 16S rRNA gene sequence phylogenies, indicate that these MAGs are all at least in novel orders within the *Thermoplasmata,* and SAL16 potentially represents a novel class within the Euryarchaeota. Here, we will refer to these as novel orders within the Thermoplasmata_A class, in agreement with the RED based taxonomic classification (Parks *et al.*, 2018), or the *Thermoplasmata* as in NCBI.

Phylogenetic trees constructed from 16 concatenated ribosomal proteins (RPs) (Figure 1), 122 proteins used by GTDB (Supplemental Figure 2), and 16S rRNA genes (Supplemental Figure 1) all show good agreement on the position of these orders as closely related to, but distinct from and basal to, the *Methanomassiliicoccales*. Some slight differences in topology are notable, however; both the RP and 122 protein-based trees show SAL16 (*Ca.* Lunaplasmata lacustris), the RBG-16 order, and the SG8-5 order MAGs as branching basal to the *Methanomassiliicoccales*, which is further supported by the 16S rRNA gene tree for the MAGs with recovered 16S rRNA genes. Placement of the order UBA10834 showed the most variation between the RP and RED-based trees. In the RP tree, UBA10834 order MAGs are grouped in a clade with the three orders listed above (though with low bootstrap support of 0.127), and this clade is basal to the *Methanomassiliicoccales* (Figure 1). However, the RED-based tree places the order UBA10834 as grouping separately from the other three orders, and more closely with the *Methanomassiliicoccales* (Supplemental Figure 2). This placement is supported by a slightly higher AAI between order UBA10834 and *Methanomassiliicoccus* MAGs than between UBA10834 MAGs and MAGs belonging to RBG-16, SG8-5, and SAL16 (*Ca*. Lunaplasma lacustris).

The highly complete (>95%) and minimally contaminated (<5%) SAL16 MAG meets the recently proposed MiMAG standards for naming organisms (Bowers *et al.*; Chuvochina *et al.*, 2019). Thus, we propose naming this microorganism *Ca.* Lunaplasma lacustris ord. nov. sp. nov. within the new order *Cand*. Lunaplasmatales (“Luna” referring to moon, “plasma” referring to being within the Ca. Thermoplasmatota, lacustris referring to lake; formal description below). This names reflects the previous enrichment of a representative of this species from carbonate cave deposits (referred to as moonmilk) in Austria (Reitschuler *et al.*, 2016; 2014), for which there is no published isolate or genomic characterization.

### Comparison of the basal lineages to *Methanomassiliicoccales*

These MAGs are deeply branching members of the same phylogenetic clade that otherwise only includes the methanogenic *Methanomassiliicoccales* (Figure 1). The cultivated *Methanomassiliicoccales* produce methane through methylotrophic methanogenesis, with variations in terms of actual and predicted substrates along with enzyme complexes involved in the metabolism of these compounds compared to more well studied methanogens (Borrel *et al.*, 2014; Speth and Orphan, 2018). A common feature of these microorganisms is a truncated methanogenesis pathway, relative to the canonical hydrogenotrophic methanogenesis pathway observed in many euryarchaeal methanogens (Borrel *et al.*, 2014; Kaster *et al.*, 2011). In the *Methanomassiliicoccales*, the transfers of methyl groups from compounds such as methylamines, methylsulfides, or methanol to Coenzyme M (CoM) to generate methyl-CoM are predicted to be mediated by methyltransferase (MtbA, MtsA/MtsB, MtaA) and methyltransferase/coronoid proteins (MtmBC, MtbBC, MttBC, MtaBC) (Borrel et al. 2013; Lang et al. 2015). The methyl group from the methyl-CoM is then reduced to methane with electrons and H^+^ supplied from H_2_. These *Methanomassiliicoccales* organisms are H_2_-dependent due to the absence of the Wood-Ljungdahl pathway that would otherwise provide reducing equivalents in a similar mechanism to conventional methylotrophic methanogens from the *Methanosarcinales* (Rother et al. 2005). Concomitantly during the reduction of methyl-CoM to methane, another enzyme called Coenzyme B (CoB) replaces the methyl group of methyl-CoM to form a heterodisulfide bond with CoM. The CoM is regenerated by the heterodisulfide/methylviologen reductase complex (HdrABC-MvhADG) with electrons generated from oxidized H_2_ in a mechanism similar to typical methanogen electron bifurcating reactions (Lang et al. 2015). The reduced ferredoxin generated from this electron bifurcation is then predicted to oxidize a membrane-bound Fpo-like + hdrD complex to drive H^+^ and/or Na^+^ translocation across the inner membrane and create the proton or sodium motive force used by ATPases to produce ATP (Borrel *et al.*, 2014; Kröninger *et al.*, 2015; Lang *et al.*, 2015). While these *Methanomassiliicoccales* genomes/MAGs contain many core methanogenesis proteins, notable exceptions include genes encoding for the N5-methyltetrahydromethanopterin:CoM methyltransferase (Mtr) complex, Fmd/FwdABCD, and methanogenesis marker genes 10 and 14 (Borrel *et al.*, 2014).

In contrast to the *Methanomassiliicoccales*, the 12 basal MAGs identified in this study lack characterized genes that would confer the potential for H_2_-dependent or other forms of methanogenesis. No *mcr* or *mcr*-like open-reading frames (ORFs) were observed in any of these MAGs (Figure 1, Supplemental Figure 5). We recognize that the lack of *mcr* gene sequences in these MAGs does not definitively exclude the possibility that such sequences were in fact present in the genomes, but were not binned within the MAGs. However, all 12 MAGs lack methanogenesis associated genes. Furthermore, our inference that these populations are not methanogenic is supported by the apparent absence of genes coding for the insertion of the amino acid pyrrolysine, along with pyrrolysine biosynthesis genes (*pylBCDS*), and these MAGs lack many of the core “methanogenesis marker genes” observed in *Methanomassiliicoccales* (Figure 1, Supplemental Figure 5) (Borrel *et al.*, 2014; Kaster *et al.*, 2011). Two exceptions were homologs of the methanogenesis marker protein 4, which was found in the three MAGs (UBA9653, RBG-13-57-23, and RBG-19FT-COMBO-56-21), and a methanogenesis marker protein 2 homolog found in RBG-13-57-23. These proteins do not have verified functions (Kaster *et al.*, 2011), so we are unable to assess the role of these putative genes in these MAGs.

The three RBG-16 and five UBA10834 order MAGs contain ORFs annotated as methyltransferases in the same protein families as found in the *Methanomassiliicoccales* (Figure 1). Most of the putative methyltransferase genes were annotated as coding for MttB (*i.e*., within the MttB superfamily) and some for the corresponding corrinoid proteins, MtbC or MttC; no putative methyltransferase alpha subunits, *e.g.* MtaA, were found to be encoded. In methanogens, MttB transfers methyl groups from trimethylamine to the corrinoid protein MttC, then MtaA or MtbA will transfer the methyl group to CoM. Many of the traditional MttB proteins in methanogens contain the amino acid pyrrolysine (Gaston *et al.*, 2011; Borrel *et al.*, 2014). However, non-pyrrolysine containing members of this protein family are widely found in non-methanogens, including MttB superfamily proteins which utilize glycine betaine (Ticak *et al.*, 2014). In the basal MAGs analyzed here, the ORFs annotated as *mttB* do not encode pyrrolysine, and they are phylogenetically distinct from *mttB* sequences of known methanogens (Supplemental figure 4), grouping instead with diverse lineages not associated with methanogens. Therefore, we infer that they encode non-methanogenic MttBs. While some of these putative *mttB* ORFs group more closely to glycine betaine reductases than the methanogenic MttBs, it is unclear what role these MttBs play in the non-methanogenic MAGs. Potentially, these MttBs may confer a methylotrophic metabolism not linked to methane formation.

Notably, ORFs annotated as *mtrAH* genes were observed in some of the environmental *Methanomassiliicoccales* MAGs, three of the SG8-5 order MAGs, but none of the other nine basal MAGs (Figure 1). In euryarchaeal methanogens a complete methyl-H_4_MPT–coenzyme-M-methyltransferase (MtrABCDEFGH) complex translocates Na^+^ ions across the cell membrane during hydrogenotrophic or acetoclastic methanogenesis, but it is not necessary for methylotrophic methanogenesis (Welander and Metcalf, 2005). Perhaps related to this, genes encoding Mtr subunits are not typically observed in the *Methanomassiliicoccales* (Borrel *et al.*, 2014). In the first observation of a *Methanomassiliicoccales* genome with *mtr* (Speth and Orphan, 2018), the authors identified *mtrAH* homologs in an environmental *Methanomassiliicoccales* MAG referred to as MAssiliicoccales Lake Pavin (MALP.) Here, we find *mtrAH* homologs in many *Methanomassiliicoccales* MAGs closely related to MALP. Both the *Methanomassiliicoccales* and the SG8-5 *mtrH* sequences group separately from other known methanogens (Supplemental Figure 3), supporting the notion that either *Methanomassiliicoccales* methanogenic abilities might have been gained through horizontal gene transfer or that the *mtrAH* genes were horizontally transferred. Speth and Orphan (2018) proposed that *mtrAH* actually encode methyltetrahydrofolate:CoM methyltransferase, which could be used in the Wood-Ljungdahl pathways found in the MALP MAG (Speth and Orphan, 2018). However, the three SG8-5 MAGs do not contain genes for the Wood-Ljungdahl pathway (Figure 3), so the *mtrAH* homologs in this lineage have an unknown function, but it is likely involved in the transfer of methyl groups.

**Figure 3.**
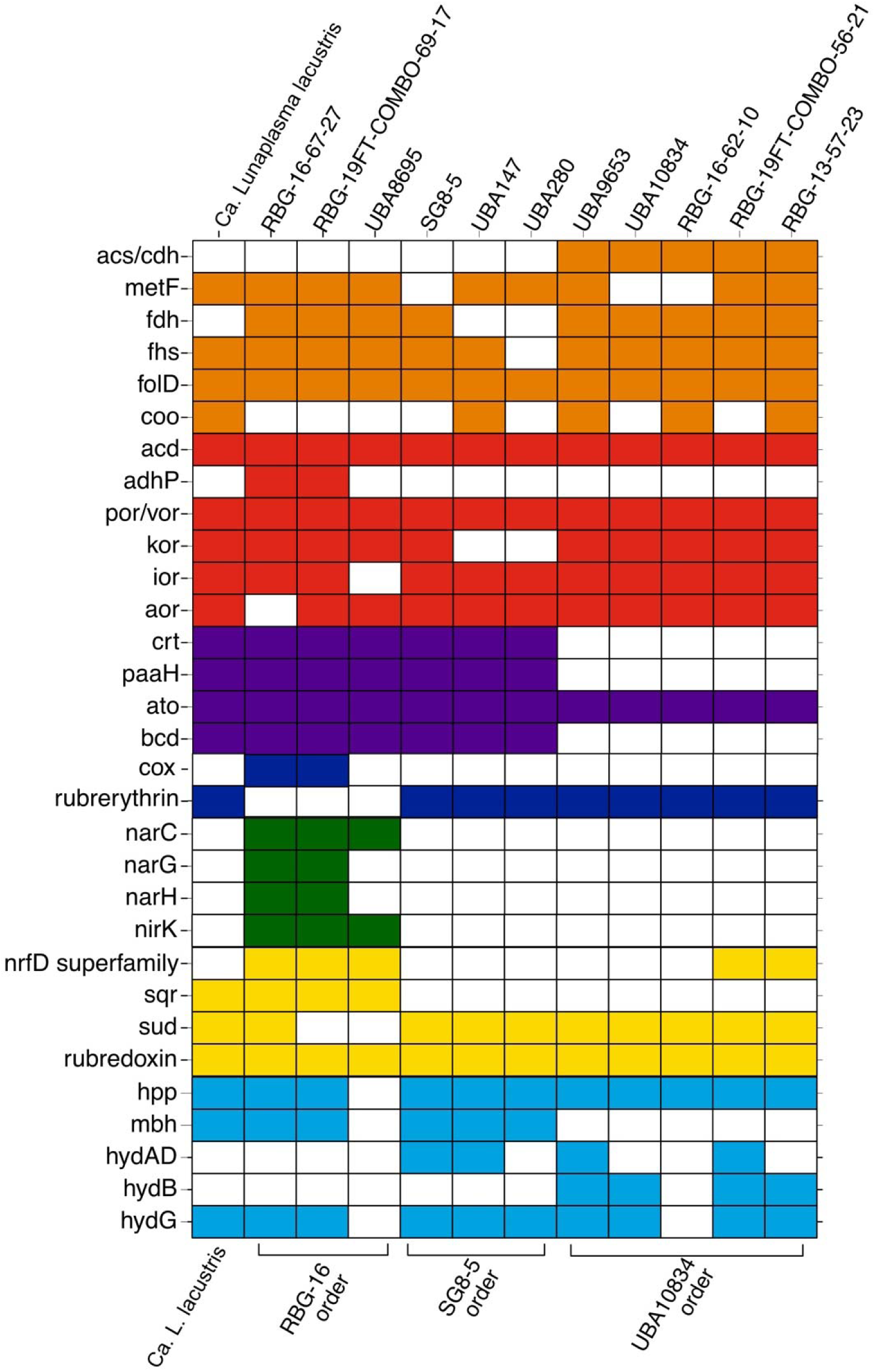
Key metabolic genes and their presence in the MAGs basal to the Methanomassiliicoccales. MAGs are grouped by their phylogenetic relationships, and putative gene assignments are grouped and colored by function or biogeochemical cycle (Orange: acetyl-CoA pathway: red: fermentation and amino acid degradation; purple: beta oxidation; dark blue: oxygen; green: nitrogen; yellow: sulfur; light blue: hydrogenases and pyrophosphatase). Gene names and products are detailed in Supplemental table 8. For genes with multiple subunits, presence was marked if at least half of the subunits were present.

### Evolutionary history of methanogenesis within the Thermoplasmata class

Taken together, these results support our conclusion that Ca. Lunaplasma lacustris, SG8-5, RBG-16, and UBA10834 order MAGs do not represent methanogenic Archaea, despite grouping phylogenetically adjacent and basal to the methanogenic *Methanomassiliicoccales*. These basal MAGs do however have some features in common with the *Methanomassiliicoccales,* such as retaining *hdrABCD* and, in some cases, a select few methanogenesis marker genes (Figure 1). Within the *Thermoplasmata*, only the *Methanomassiliicoccales* are known to be methanogenic, with orders including the *Thermoplasmatales* and the uncultured Candidatus Poseidoniales (previously known as Marine Group II or MGII) containing no known methanogens or methanotrophs (Tully, 2019; Rinke *et al.*, 2019).

It has previously been proposed that methanogenesis was lost in the *Thermoplasmatales* and retained in the *Methanomassiliicoccales* (Evans *et al.*, 2019), but with the addition of extra genomes here, it now appears that the *Methanomassiliicoccales*-related MAGs could have gained genes for methane metabolism, further complicating the evolutionary history of metabolic properties in the *Thermoplasmata*. Given the paraphyletic nature of groups basal to the *Methanomassiliicoccales* within the *Thermoplasmata* (Figure 1), the most parsimonious explanation for the lack of methanogenesis in these basal groups is that methanogenesis was in fact not present in the last common ancestor of the *Thermoplasmata.* Instead, methanogenesis was gained by an ancestor of the *Methanomassiliicoccales* through horizontal gene transfer, as hypothesized for the *Ca.* Bathyarchaeota BA1 and BA2 and *Ca.* Verstraetearchaota lineages (Evans *et al.*, 2015; Vanwonterghem *et al.*, 2016). Although the phylogenetic trees presented here show topological differences (including two clades vs. one clade of these MAGs basal to the *Methanomassiliicoccales*, as described above), the preservation of vertical transmission of methanogenesis within the *Thermoplasmata* class to the *Methanomassiliicoccales* would still require more than one loss of methanogenesis in the *Thermoplasmata*. However, as *mcr* or *mcr*-like genes are found throughout the Archaeal tree, potentially mostly due to vertical descent (Hua *et al.*, 2019), then the less parsimonious explanation (an inference of multiple losses of methanogenesis in the *Thermoplasmatales* and other lineages basal to *Methanomassiliicoccales*) is entirely possible.

### Metabolic potential of lineages basal to the *Methanomassiliicoccales*

The *Ca.* Lunaplasma lacustris MAG was the highest quality MAG in our dataset, thus we focused our metabolic analyses on this lineage, examining ORF annotations to attribute potential metabolic functional to this MAG. For the less complete MAGs in the SG8-5, UBA10834, and RBG-16 orders, we examined the MAGs as groups and compared these predicted functions to those of *Ca*. Lunaplasma lacustris. In all cases, these MAGs were previously either metabolically uncharacterized (MAGs beginning with the prefix “UBA” (Parks *et al.*, 2018) or characterized as part of larger studies of many MAGs (SG8-5 as in (Lazar *et al.*, 2017) MAGs beginning with the prefix “RBG” (Anantharaman *et al.*, 2016)).

Annotation suggests that *Ca.* Lunaplasma lacustris likely can conserve energy through amino acid metabolism, similar to the metabolism proposed for the MBG-D Single Amplified Genome (SAG) in (Lloyd *et al.*, 2013). In *Ca*. Lunaplasma lacustris, amino acids and short peptides are predicted to be imported into the cell by branched chain and polar amino acid transporters (Figure 2). Aminotransferases are predicted to convert the amino acids into 2-keto acids, which could be oxidized through the action of the oxidoreductases Ior, Vor, or Kor. Ferredoxin reduction is predicted to be coupled to amino acid oxidation (Sapra *et al.*, 2003; Lloyd *et al.*, 2013). Ferredoxin (Fd_red_) can then be oxidized by a membrane-bound Fpo-like + HdrD and/or glcD complex and a membrane-bound hydrogenase, creating a proton motive force. ATP can then be produced through a V/A-type ATPase.

**Figure 2.**
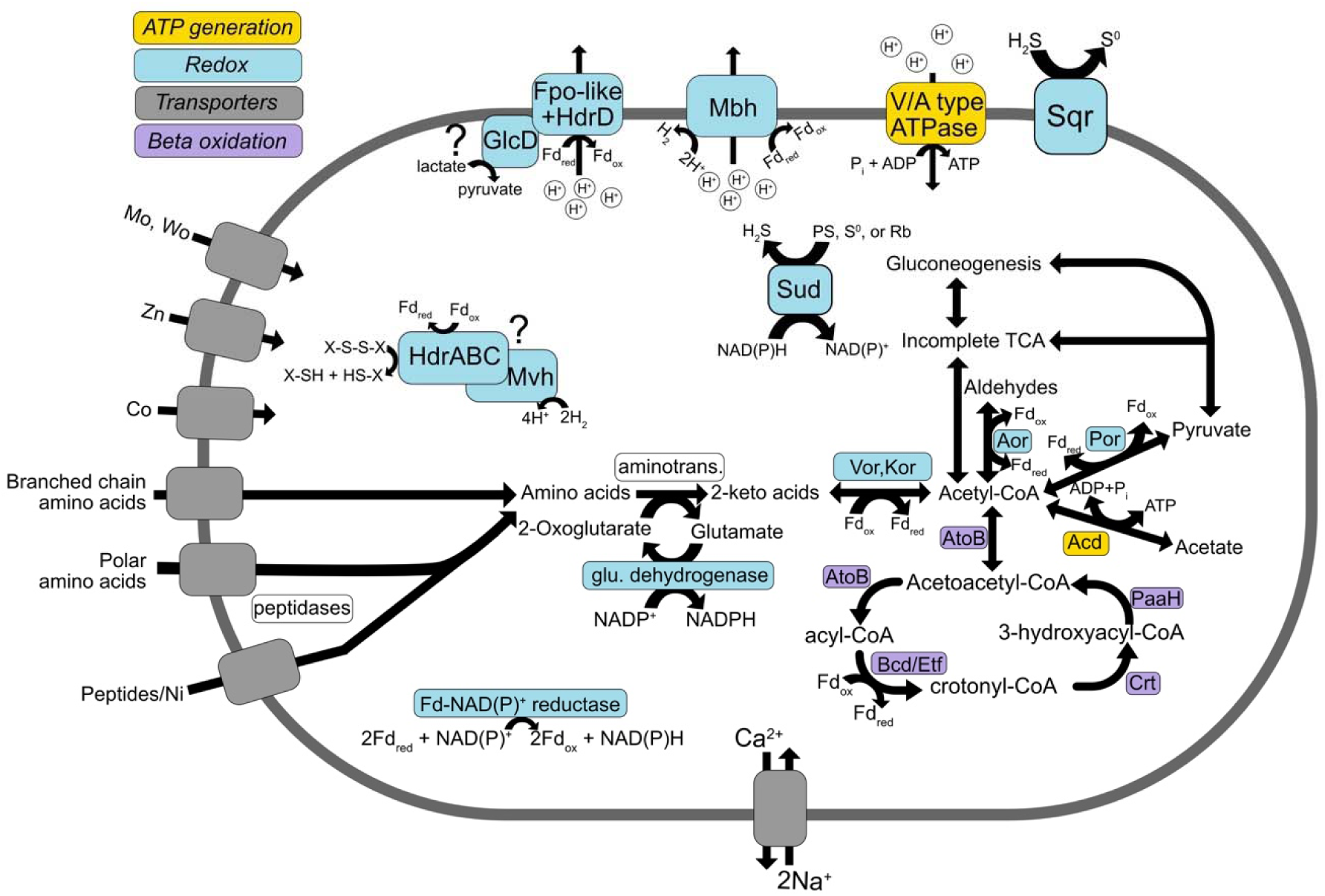
Proposed metabolic scheme for *Ca*. Lunaplasma lacustris based on genome reconstruction. *Ca*. L. lacustris putatively uptakes peptides or amino acids through transporters, then intracellular peptidases break down peptides. Aminotransferases and glutamate dehydrogenases convert amino acids into 2-keto acids, which can be oxidized to acetyl-CoA by various oxidoreductases (Ior, Vor, Kor), producing reduced ferredoxin (Fd_red_). Oxidation of acetyl-CoA by Aor into aldehyldes could serve as another source of Fd_red_. Fpo-like subunits complexed with HdrD could drive a proton (or sodium) motive force using reducing power from Fd_red_. A membrane-bound hydrogenase similarly could perform this reaction, coupling Fd oxidation to hydrogen production. A beta-oxidation pathway similar to that proposed in the *Ca*. Polytropus marinifundus, or substrate-level oxidation through Acd could be used for additional reducing power or ATP synthesis. ORFs annotated as sulfide:quinone reductase (Sqr) and sulfide dehydrogenase (SuDH) genes suggest sulfur compounds as a source of energy as well, potentially including sulfide oxidation or S_0_ or polysulfide (PS) oxidation.

Multiple ORFs with homology to hdrABC/mvhADG subunits were found in *Ca.* Lunaplasma lacustris (Figure 2; Supplemental Table 9). In some methanogens, the HdrABC/MvhADG complex reduces the heterodisulphide CoM-S-S-CoB and Fd_red_ (Buan and Metcalf, 2010). However, Hdr subunits are also found in non-methanogenic sulfate-reducing bacteria and archaea, where they have been proposed to function in electron bifurcation associated with sulfur metabolism pathways (Ramos *et al.*, 2015). In *Ca.* Lunaplasma lacustris, this complex could be coupling hydrogen oxidation to the reduction of ferredoxin and an unknown heterodisulphide or other sulfur compound, as proposed for SG8-5 (Lazar *et al.*, 2017), *Archaeoglobus profundus* (Mander *et al.*, 2004), and an uncultured bacterial population (Castelle *et al.*, 2013).

A beta-oxidation pathway showing similarity, based on the presence of genes similar to *atoB, paaH, crt,* and *bcd/etf*, to those described in the methanogenic *Archaeoglobus* was also observed in *Ca.* Lunaplasma lacustris (Boyd *et al.*, 2019), so it is possible that these organisms degrade fatty acids for energy conservation (Figure 2; Supplemental Table 8). ORFs annotated as acetate-CoA ligase (*acd)* were also present, possibly conferring the ability to produce ATP through fermentation (Schäfer *et al.*, 1993) and providing another route for energy conservation in this apparently versatile population.

The three RBG16 order MAGs (RBG-19FT-COMBO-69-17, RBG-16-67-27, and UBA8695; Supplemental Table 10-12) are the most closely related to *Ca.* Lunaplasma lacustris, and they appear to have similar metabolic potential, *i.e.*, amino acid respiration, fermentation, and beta oxidation (Figure 3). These MAGs and *Ca.* Lunaplasma lacustris also contain ORFs annotated as a sulfide-quinone reductase (*sqr*). In some organisms, Sqr can oxidize H_2_S to S_(n)_, which can be coupled to oxygen reduction (Brito *et al.*, 2009; Lencina *et al.*, 2013). In addition to putative *sqr* genes, these MAGs encode rubredoxin, an electron carrier found in some sulfur respiring organisms (Ma *et al.*, 1993; Lumppio *et al.*, 2001), and sulfur dehydrogenases, which can act as bifunctional S^0^ or S_n_ reductases or Fd:NADPH oxidoreductases (Ma and Adams, 1994). Based on the presence of these putative genes, it is likely that these MAGs can participate in sulfur cycling by consuming or producing sulfides.

A distinctive difference between the RBG-16 cluster MAGs and *Ca.* Lunaplasma lacustris is the presence of ORFs annotated as archaeal nitrate reductase complex (*narGH*) (Yoshimatsu *et al.*, 2000) and nitrite reductase *nirK* in RBG-16-67-27 and RBG-19FT-COMBO-69-17 (Fig. 3). In the less complete UBA8695 (70.76% estimated completeness), *narGH* homologs were not found. UBA8695 did however contain a putative *narC*, which encodes cytochrome b-561 in the archaeon *Haloarcula marismortui* (Yoshimatsu *et al.*, 2007). This indicates that these MAGs likely represent nitrate- and nitrite-reducing archaea, which is consistent with their brief metabolic characterization in the source publication, which examined key functional genes in over 2,500 genomes (Anantharaman *et al.*, 2016). Two MAGs in the RBG-16 order contain cytochrome C oxidase subunits, pointing to the capacity for aerobic respiration (Capaldi *et al.*, 1983). Lending further support to this is the likely lack of rubrethythrin, a protein used to combat oxidative stresses in anaerobes, in the RBG-16 order (Weinberg *et al.*, 2004; Lencina *et al.*, 2013).

ORFs annotated as part of the *nrfD* family, which can function in nitrite oxidation and sulfur/polysulfide reduction (Hussain *et al.*, 1994), were also found in the RBG-16 order and two UBA10834 group MAGS (RBG-19FT-COMBO-56-21 and RBG-13-57-23) (Figure 3; Supplemental Tables 16 and 17). The putative *nrfD* subunits in these MAGs were all located next to a *dmsB* gene (Supplemental Figure 6), which can encode part of a dimethyl sulfoxide or trimethylamine N-oxide reductase complex (Müller and DasSarma, 2005). Only one MAG, RBG-19FT-COMBO-56-21, also contained *dmsA* and *dmsD* in the same operon as *dmsB* and a *nfrD* subunit, which was alternatively annotated as *dmsC*. In RBG-13-57-23, *nrfD* was also located next to a formate dehydrogenase subunit (*fdhD*), and in RBG-19FT-COMBO-69-17, one *nrfD* was between a molybdopterin oxidoreductase encoding gene annotated as encoding tetrothionate reductase subunit A (*ttrA)* and *dmsB*. Boyd et al. (2019) found *dmsB* and *nrfD* located on the same operon in an Archaeoglobus MAG recovered from the deep subseafloor and posited that these genes encode proteins used in sulfur redox chemistry (Boyd *et al.*, 2019). Similarly, sulfate-reducing organisms use NrfD-like proteins as electron shuttles during sulfate reduction (Pereira *et al.*, 2011). These ORFs indicate that the reduction of sulfur compounds (S^0^, PS, and/or DMSO) may be a metabolic strategy in these organisms, but due to the various functionalities of the predicted gene products, it remains inconclusive exactly which metabolisms they confer.

SG8-5 order MAGs also appear to have a similar metabolic scheme to *Ca.* Lunaplasma lacustris, with some notable deviations and variability among genomes (Figure 3; Supplemental Tables 13-15). In Lazar *et al.*, 2017, SG8-5 was described as a peptide degrader, which supports our findings that all three SG8-5 order organisms could be peptide-degrading. A difference between the three MAGs, though, is that SG8-5 was hypothesized to be using HdrABC/MvhADG as part of its machinery to reduce ferredoxin Lazar *et al.*, 2017. However, UBA147, which is 89.6% complete compared to SG8-5’s 72.24% completeness, does not contain any predicted *hdr* genes, nor does UBA280 (77.8% completeness) (Figure 1). This suggests that Hdr may not play a role in UBA147, and instead the membrane-bound hydrogenases and potentially the P-type pyrophosphatases are presumed to be the main drivers of the proton motive force.

The predicted metabolic schemes in the order UBA10834 are perhaps the most divergent from the other MAGs basal to the *Methanomassiliicoccales*. Though these MAGs still contain ORFs indicative of amino acid respiration and fermentation as likely metabolic strategies, these MAGs lack evidence of beta-oxidation pathways (Figure 3). However, they contain ORFs annotated as acetyl-CoA synthase/carbon monoxide dehydrogenase, which along with those for the Wood-Ljungdahl pathway are required for carbon fixation (Ragsdale and Pierce, 2008). Several MAGs in the UBA10834 order contained multiple ORFs annotated as sulfhydrogenase (*hydABDG)*, which can function as a hydrogenase and S^0^ or S_n_ reductase (Ma *et al.*, 1993), though the function here is not confirmed. Thus the UBA10834 MAGs may represent autotrophic organisms that can couple sulfur reduction to hydrogen oxidation.

### Global distribution of populations basal to the *Methanomassiliicoccales*

The NCBI non-redundant nucleotide (nt) database was searched to determine where 16S rRNA gene sequences from these *Methanomassiliicoccales*-related populations have been previously observed. 16S rRNA gene sequences recovered from three MAGs were used for this analysis (Supplemental Table 4). The 16S rRNA gene sequences from UBA147 and SG8-5 were found in globally distributed samples from environments such as sediments, coal beds, oil-sands tailing ponds, and groundwater (Fig. 4; Supplemental Table 21). These findings are consistent with the environmental origins of the MAGs recovered from these lineages and suggest that these lineages may be exclusively environmental. This is in contrast to some *Methanomassiliicoccales* lineages that, thus far, have been exclusively found in the human and animal gastrointestinal tracts (Figure 1) (Söllinger *et al.*, 2015).

**Figure 4.**
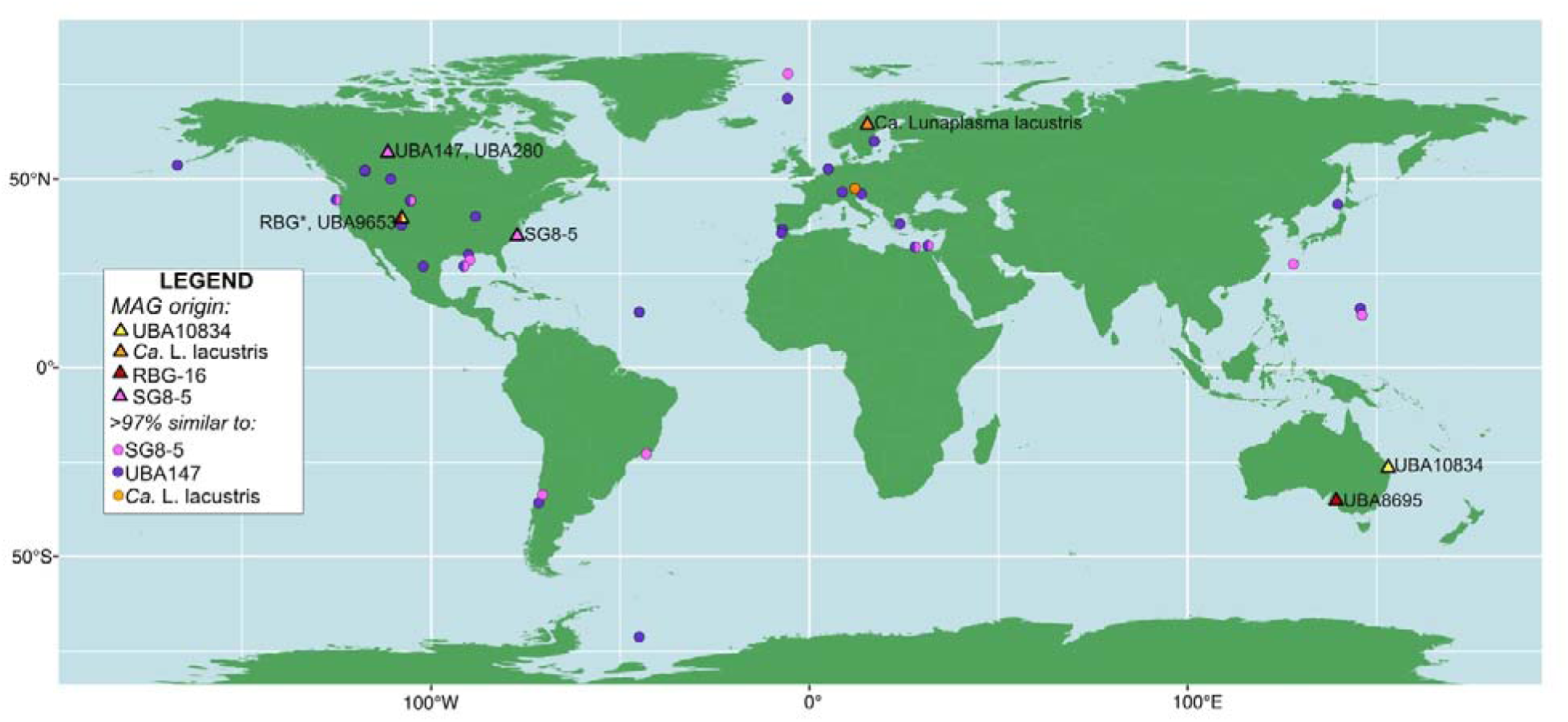
Map of MAGs (triangles) and 16S rRNA gene fragments (circles) in the NCBI nucleotide (nt) database within 97% similarity to 16S rRNA genes recovered in three MAGs from two orders: SG8-5 (SG8-5 and UBA147) and Lunaplasmatales (*Ca*. L. lacustris) (16S rRNA gene sequences were not recovered from order RBG-16 or UBA10834 MAGs). All MAGs and sequences were from environmental samples (that is, not host-associated). Shapes with multiple colors indicate recovery of multiple orders from the same site. MAG IDs are next to their locations. *The five MAGs beginning with “RBG” were binned from data generated from a Rifle creek site in Colorado, United States.

The environments containing 16S rRNA gene sequences similar to UBA147 and SG8-5 tended to be anoxic, which is consistent with this lineage having no evidence of aerobic metabolism, unlike the RBG-16 order lineages discussed here. These anaerobic environments included methane-rich environments, such as the methanogenic coal beds in the Powder River Basin in the western United States and methane cold seeps in the northeast Pacific Ocean (Barnhart *et al.*, 2013; Marlow *et al.*, 2014). The majority of the UBA147-related sequences were from anaerobic zones of tailing ponds in Alberta, Canada. The SG8-5 related sequences were found in similar, and sometimes the same, environments as UBA147-related sequences (Figure 4). However, apart from the coalbed-associated sequences, the SG8-5-related sequences were recovered from marine influenced areas, such as bays, continental margins, or deep-sea sediments, and the SG8-5 MAG itself was recovered from estuary sediments. It thus appears that within the order SG8-5, UBA147 represents a more widely distributed clade, whereas SG8-5-like organisms tend to be more marine-associated.

In contrast to the ubiquitous UBA147 and SG8-5 sequences, 16S rRNA sequences from approximately the same species as *Ca.* Lunaplasma lacustris (i.e. defined as > 97% similar) were found exclusively in calcite deposits (‘moonmilk deposits’) from alpine caves in the Austrian Alps (Reitschuler et al, 2016). Intriguingly, Reitschuler *et al*. were able to enrich these closely related ‘Moonmilk Archaea’ under anaerobic conditions at low temperatures (10°C), concluding that these organisms were unlikely to be methanogenic, methanotrophic, or nitrate- or iron-reducing. These conclusions support our findings that *Ca.* Lunaplasma lacustris is likely to be heterotrophic and non-methanogenic. As the *Ca*. Lunaplasma lacustris MAG was recovered from methanogenic sediments in post-glacial freshwater lakes that have seasonal freeze-thaw cycles in northern Sweden (Seitz *et al.*, 2016), it is possible that these organisms prefer cold, anoxic environments, consistent with the metabolic predictions for this lineage.

## Description of Ca. Lunaplasma lacustris

*Candidatus* Lunaplasma (Lu.na.plas’ma. N.L. fem. n. *luna* (from Latin. fem. noun. *luna*) moon, Gr. neut. n. *plasma*, something formed or molded, a form). *Candidatus* Lunaplasma lacustris (la.cus.tris. M.L. fem. adj., lacustris inhabiting lake). “Lunaplasmatales” (Lu.na.plas’ma.tal.es N.L. n. “Lunaplasmatales” -entis, type genus of the family; suff. -ales, ending to denote an order; N.L. neut. pl. n. Lunaplasmatales, the order of the genus “Lunaplasma”).

## Conclusions

From our results we find that four orders of uncharacterized archaea closely related to *Methanomassiliicoccales* methanogens do not have pathways that would suggest a methanogenic lifestyle. This result is inconsistent with NCBI and RDP database classifications of these MAGs as belonging to the methanogenic order *Methanomassiliicoccales*. Instead, MAGs in these orders (basal to *Methanomassiliicoccales*) contain genes potentially conferring various combinations of sulfur, nitrogen, hydrogen, and aerobic metabolisms, with one order possibly autotrophic. Relationships of 16S rRNA gene sequences to those from published amplicon datasets indicate that species in one order, SG8-5, are widespread in both marine and terrestrial environments, while the *Ca*. Lunaplasmata lacustris MAG apparently represents a species with a constrained ecological distribution. In combination with phylogenetic analyses of the *Methanomassiliicoccales* and its basal lineages, our metabolic reconstructions suggest an alternative evolutionary history of methanogenesis in the *Thermoplasmata*, relative to a recent hypothesis that suggested that methanogenesis was present in the ancestor of the *Thermoplasmata* and vertically transmitted to the *Methanomassiliicoccales* but lost in the *Thermoplasmatales*. We suggest instead that the capacity for methanogenesis was absent in the ancestor of the *Thermoplasmata* and was gained in the ancestor of the *Methanomassiliicoccales* via horizontal gene transfer after divergence from the basal lineages analyzed here. Another possible explanation is that the ability to perform methanogenesis was present in a basal ancestor to the *Thermoplasmatales,* but was lost multiple times in the descending lineages. These findings highlight the potential roles of non-methanogenic *Thermoplasmata* in the environment and contribute to our rapidly developing understanding of the evolution of methanogenesis.

## Acknowledgements

We thank Elaina Graham and Lily Momper for advice on phylogenetic tree construction, and members of the NSF funded Northern Ecosystems Research for Undergraduates REU Site (EAR#1063037) team Kaitlyn Steele, Martin Wik, and Nancy Freitas for sample collection. The support and resources from the High Performance Computing group in the Bioinformatics Core at the University of California, Davis are gratefully acknowledged. Our funding sources include new laboratory start-up to JBE from the UC Davis College of Agricultural and Environmental Sciences and the UC Davis Department of Plant Pathology, and SAL-16 MAG recovery via support from the Genomic Science Program of the United States Department of Energy Office of Biological and Environmental Research, grants DE-SC0010580 and DE-SC0016440 to VIR and GWT.

---

**Supplemental figure 1.**
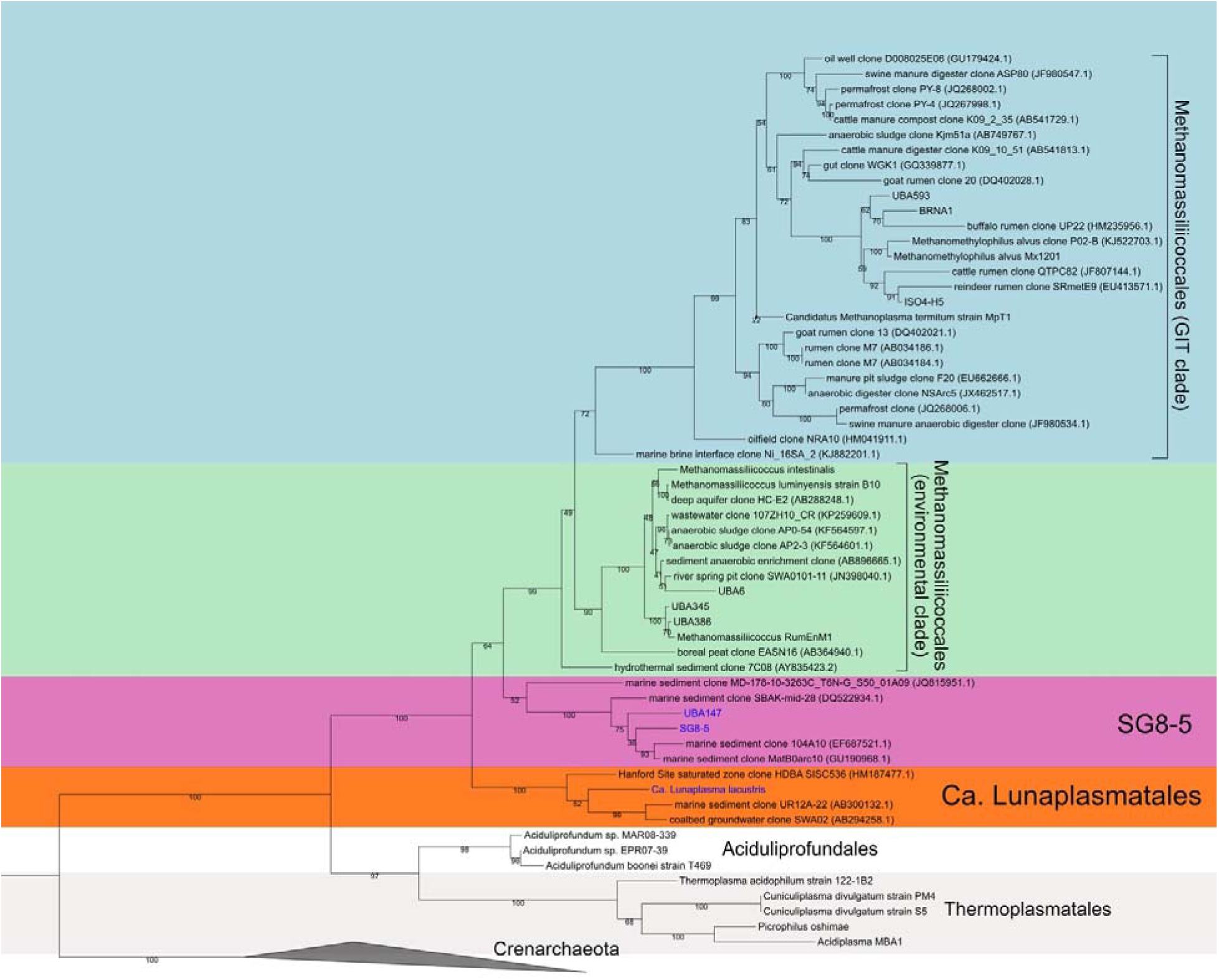
16S rRNA gene phylogenetic tree, including the three 16S rRNA gene sequences recovered from three MAGs in two orders basal to the *Methanomassiliicoccales* (blue text): Lunaplasmatales (*Ca*. Lunaplasma lacustris) and SG8-5 (UBA147 and SG8-5). The Crenarchaeota were used as an outgroup. Clades of *Methanomassiliicoccales* are labeled as in Soellinger et al. (2016). Bootstrap values are calculated from 1000 bootstraps. Background colors are as in Figure 1.

**Supplemental figure 2.**
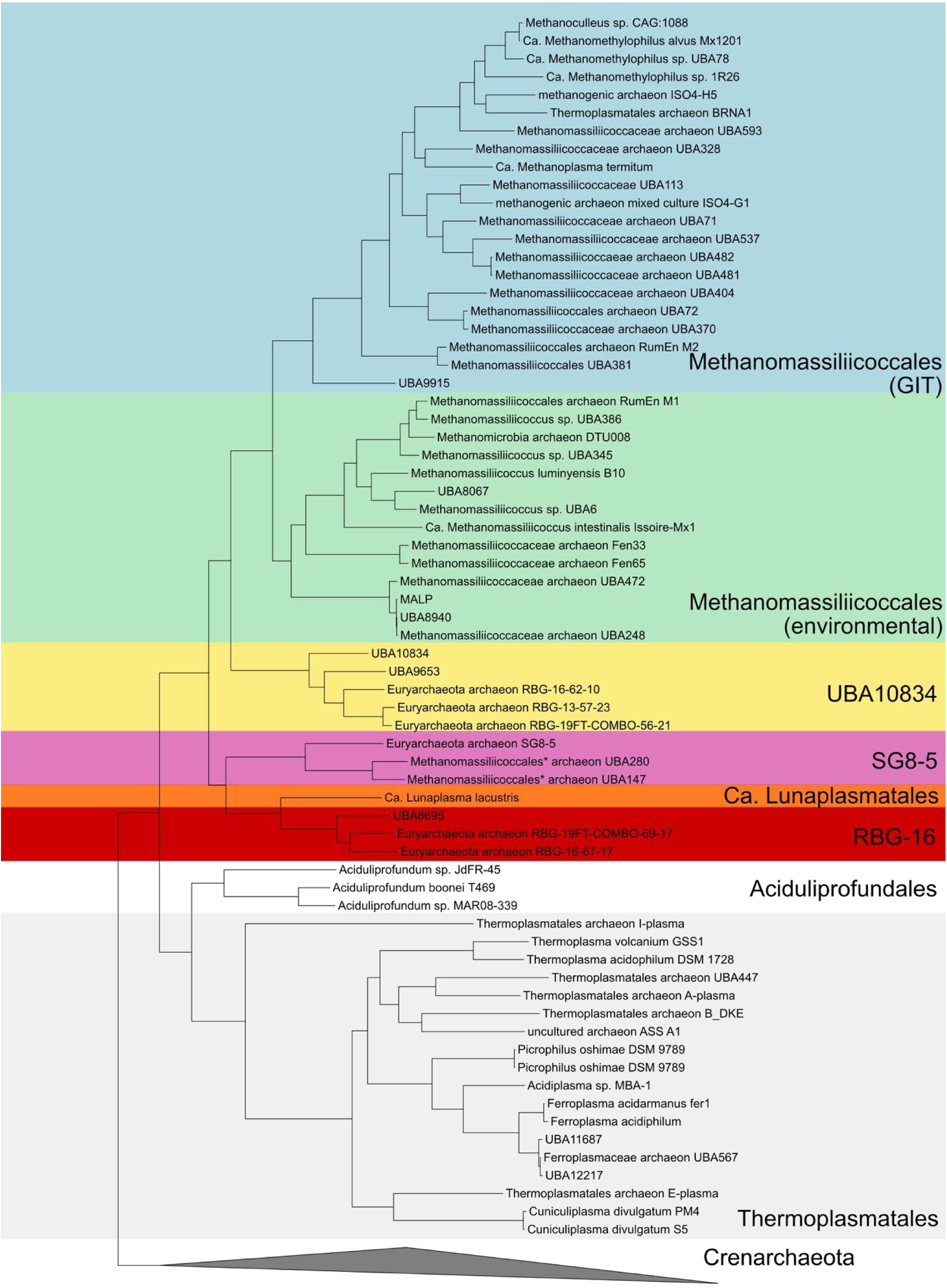
Phylogenetic tree of MAGs and genomes, based on relative evolutionary divergence (RED), as calculated by the Genome Taxonomy Database toolkit (GTDB-Tk). Crenarchaeota were used as the outgroup.

**Supplemental figure 3.**
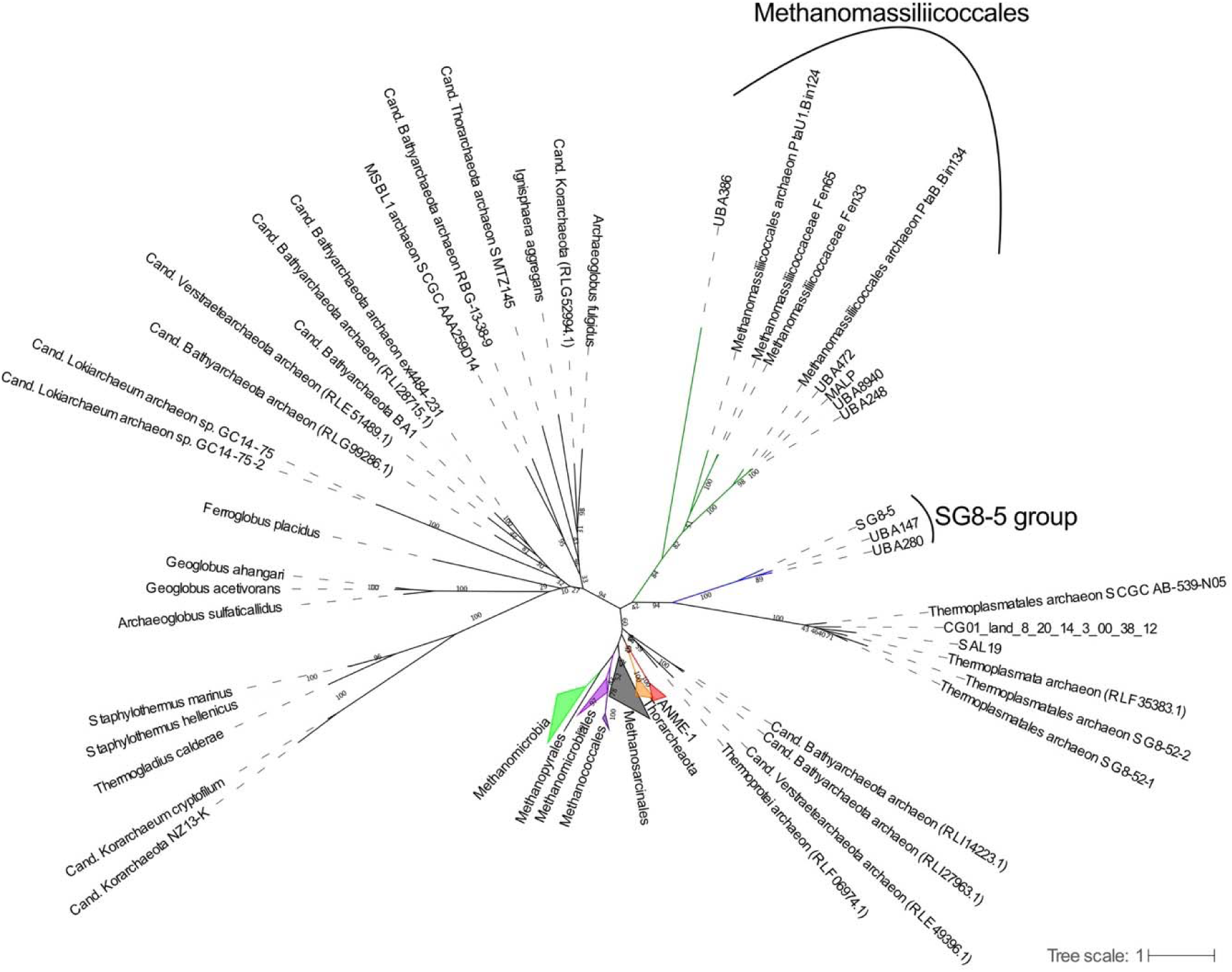
Tree of MtrH subunits, including those from *Methanomassiliicoccales* and the SG8-5 order. Branches are colored to highlight various clades. Bootstrap support (out of 100 bootstraps) is shown along branches. Dashed lines extending branches do not represent branch lengths, and are present to connect branch labels to branches for easier reading.

**Supplemental figure 4.**
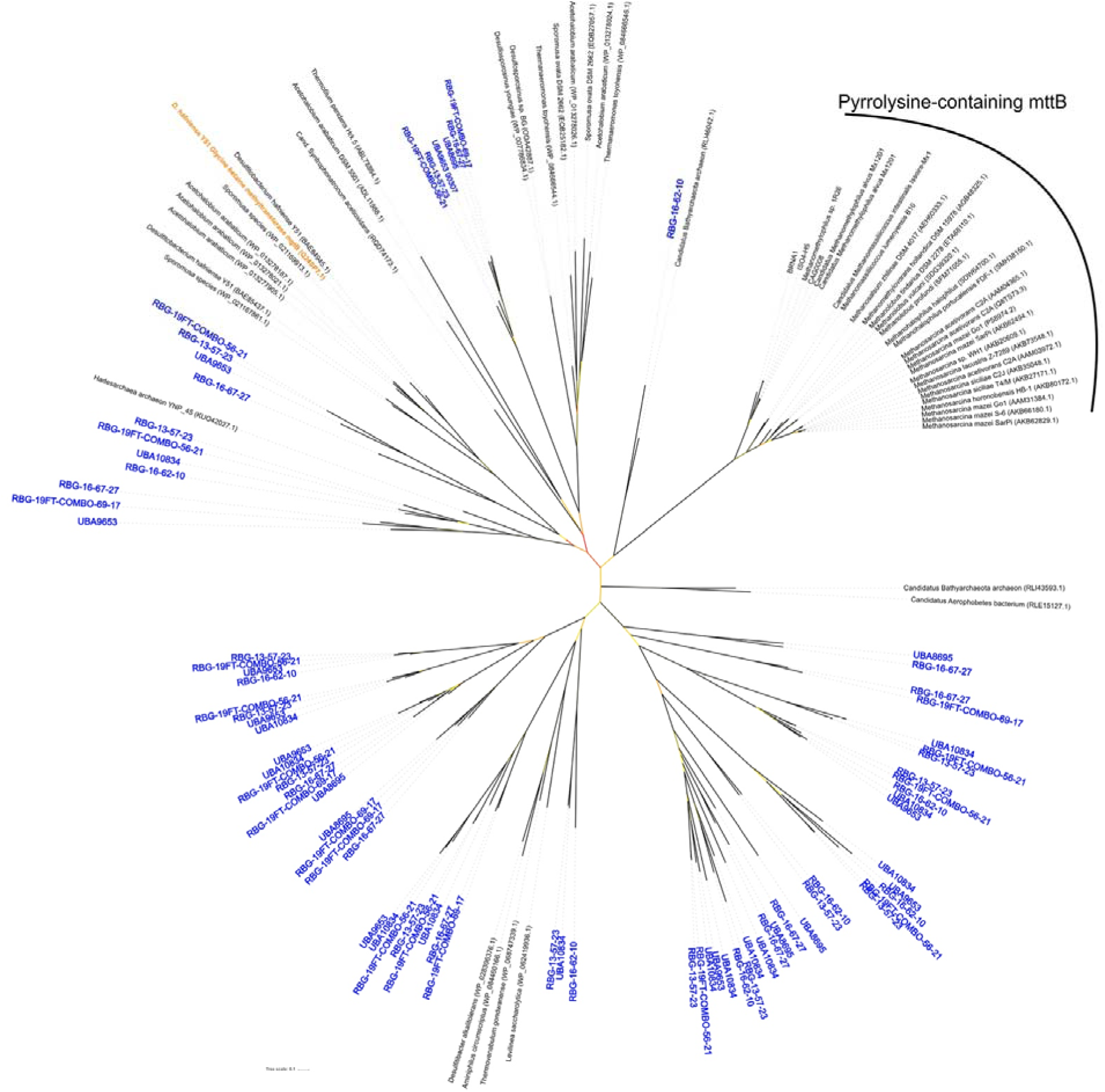
MttB superfamily tree including MttB sequences from euryarchaeal methanogens and *Methanomassiliicoccales*, glycine betaine transferases (part of the MttB superfamily but functionally distinct; in orange), and predicted MttB-like sequences recovered the MAGs basal to the *Methanomassiliicoccales* (all blue labels). Bootstrap support (out of 100 bootstraps) is indicated by branch color, displayed as a gradient, where black is the highest support (up to 100), red is the lowest support (5 or more), yellow is the midpoint (52). Dashed lines extending branches do not represent branch lengths, and are present to connect branch labels to branches for easier reading.

**Supplemental figure 5.**
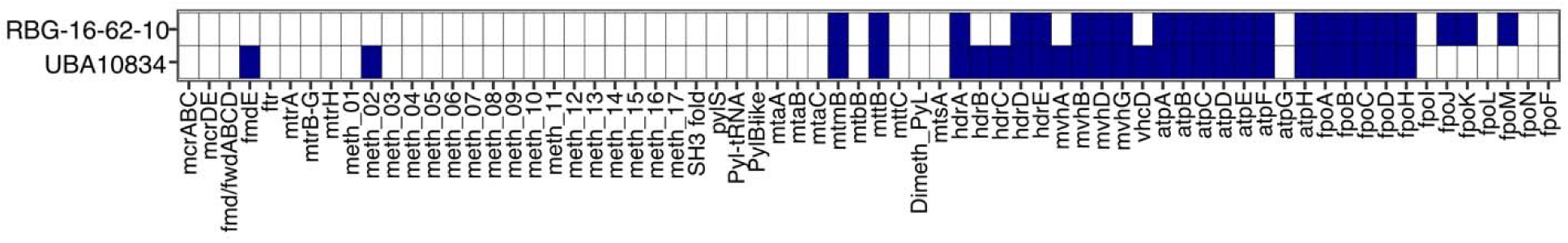
Presence of methanogenesis-related ORFs as in Figure 1 for two UBA10834 order MAGs that did not have the minimum 8 of 16 RP required for inclusion in the RP tree.

**Supplemental Figure 6.**
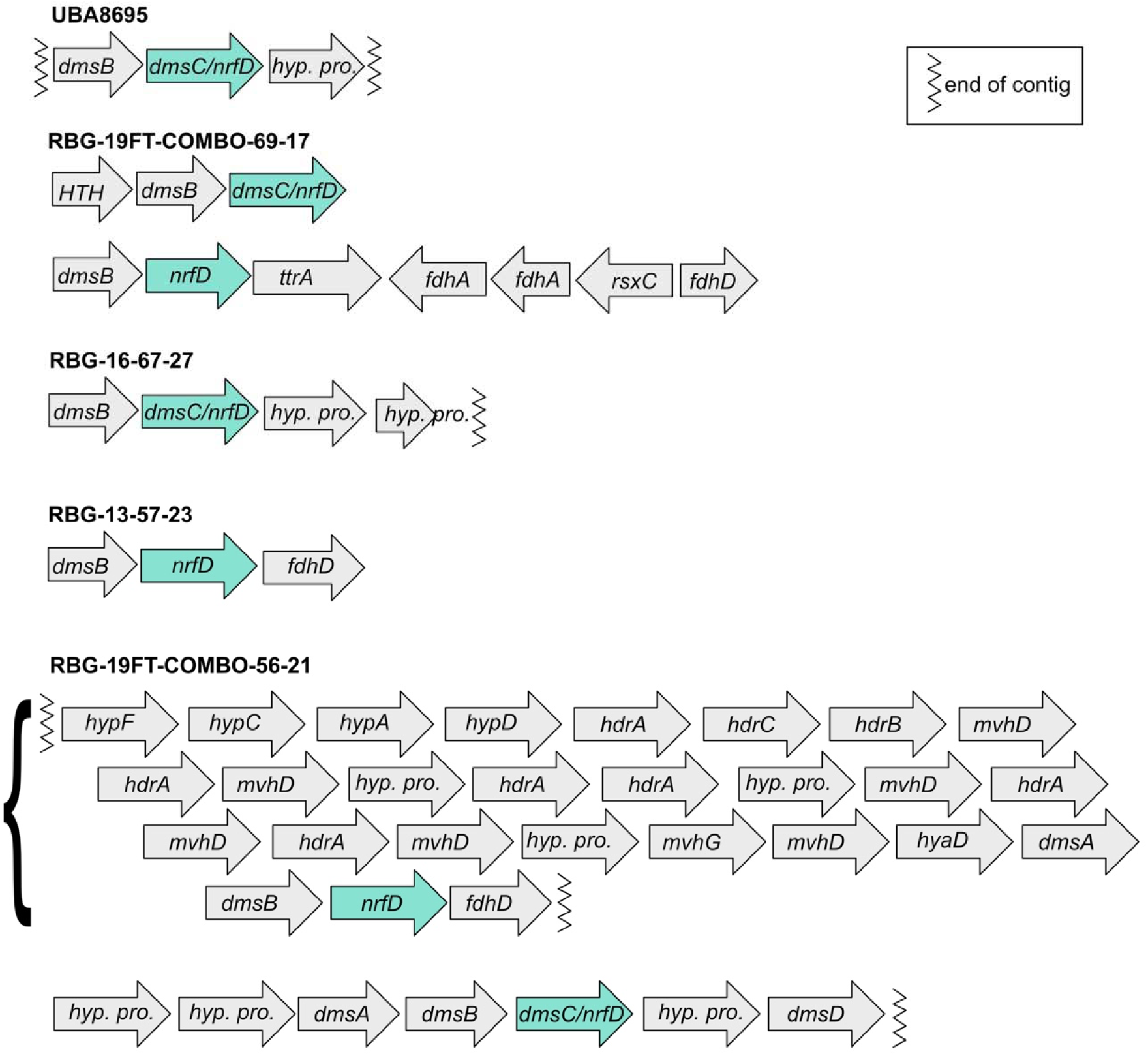
Operons containing *nrfD* annotated genes from MAGs basal to the *Methanomassiliicoccales*. Ends of contigs are marked with the zipper line. The bracket to the left of the top RBG-19FT-COMBO-56-21 contig indicates that the four lines of arrows are putatively one operon (*i.e.*, the four lines should be read as one continuous line).

Supplemental table 1. Sources of genomes and MAGs used in this study. SRA source for MAGs retrieved from GTDB are from the source metagenomes of the MAGs.

Supplemental table 2. Detailed genome stats from CheckM for all genomes and MAGs.

Supplemental table 3. Genome statistics calculated by the MiGA webserver.

Supplemental table 4. Taxonomic classification of 16S rRNA gene sequences from UBA147, SG8-5, and *Ca*. Lunaplasma lacustris by RDP classifier, Silva SINA, and Blastn searches.

Supplemental table 5. Taxonomic classification and novelty of MAGs by MiGA and GTDBTk.

Supplemental table 6. MiGA calculated AAI comparison to genomes/MAGs in the NCBI prokaryotic database.

Supplemental table 7. AAI between MAGs and genomes here, computed by the envi-omics AAI calculator.

Supplemental table 8. Genes used in Figures 1, 2, and 3.

Supplemental table 9. Comparisons of three 16S rRNA gene sequences recovered from metagenome-assembled genomes (MAGs) in this study with 16S rRNA gene sequences recovered from previous amplicon studies.

Supplemental table 10. Functional annotations from prokka, BlastKOALA, and InterProScan for *Ca.* L. lacustris.

Supplemental table 11. Functional annotations from prokka, BlastKOALA, and InterProScan for RBG-19FT-COMBO-69-17.

Supplemental table 12. Functional annotations from prokka, BlastKOALA, and InterProScan for RBG-16-67-27.

Supplemental table 13. Functional annotations from prokka, BlastKOALA, and InterProScan for UBA8695.

Supplemental table 14. Functional annotations from prokka, BlastKOALA, and InterProScan for SG8-5.

Supplemental table 15. Functional annotations from prokka, BlastKOALA, and InterProScan for UBA147.

Supplemental table 16. Functional annotations from prokka, BlastKOALA, and InterProScan for UBA280.

Supplemental table 17. Functional annotations from prokka, BlastKOALA, and InterProScan for RBG-19FT-COMBO-56-21.

Supplemental table 18. Functional annotations from prokka, BlastKOALA, and InterProScan for RBG-13-57-21.

Supplemental table 19. Functional annotations from prokka, BlastKOALA, and InterProScan for RBG-16-62-10.

Supplemental table 20. Functional annotations from prokka, BlastKOALA, and InterProScan for UBA9653.

Supplemental table 21. Functional annotations from prokka, BlastKOALA, and InterProScan for UBA10834.

